# Neural basis of melodic learning explains cross-cultural regularities in musical scales

**DOI:** 10.1101/2022.11.01.512632

**Authors:** Claire Pelofi, Mohsen Rezaeizadeh, Morwaread M. Farbood, Shihab Shamma

## Abstract

**Summary:** Seeking exposure to unfamiliar experiences constitutes an essential aspect of the human condition, and the brain must adapt to the constantly changing environment by learning the evolving statistical patterns emerging from it. Cultures are shaped by norms and conventions and therefore novel exposure to an unfamiliar culture induces a type of learning that is often described as implicit: when exposed to a set of stimuli constrained by unspoken rules, cognitive systems must rapidly build a mental representation of the underlying grammar. Music offers a unique opportunity to investigate this implicit statistical learning, as sequences of tones forming melodies exhibit structural properties learned by listeners during short- and long-term exposure. Understanding which specific structural properties of music enhance learning in naturalistic learning conditions reveals hard-wired properties of cognitive systems while elucidating the prevalence of these features across cultural variations. Here we provide behavioral and neural evidence that the prevalence of non-uniform musical scales may be explained by their facilitating effects on melodic learning. In this study, melodies were generated using an artificial grammar with either a uniform (rare) or non-uniform (prevalent) scale. After a short exposure phase, listeners had to detect ungrammatical new melodies while their EEG responses were recorded. Listeners’ performance on the task suggested that the extent of statistical learning during music listening depended on the musical scale context: non-uniform scales yielded better syntactic learning. This behavioral effect was mirrored by enhanced encoding of musical syntax in the context of non-uniform scales, which further suggests that their prevalence stems from fundamental properties of learning.

## Results and discussion

Music—like language—is produced in all known human cultures [1; 2; 3] and displays varying structural norms across cultures [4; 5; 6; 7] as well as some structural stability [8; 9; 10]. The balance between diversity and stability in cross-cultural musical structure may be governed by general principles of cultural evolution [11; 12; 13], although there is still limited empirical evidence supporting this hypothesis [14; 15]. Yet, gaining insight into the way universal structures are selected may shed considerable light on the underlying cognitive processes [16; 15; 17; 14]. This study is dedicated to examining the cause behind the prevalence of a specific musical feature, the non-uniformity of musical scales, in light of its potential role in promoting statistical learning of melodies (given that scales directly affect the way the pitches of notes are organized to form melodies). Cross-cultural studies have demonstrated that listeners’ responses to music reflect their own long-term cultural exposure [18; 19; 20; 21; 22; 7; 23; 24]. These are echoed by other studies reporting similar results but after only short-term exposure to an unfamiliar musical system [15; 25; 26; 27; 28]. In either case, this line of research has demonstrated that upon musical exposure, listeners implicitly acquire the structural properties of their exposure corpus [29; 30; 31]. A recent behavioral study provided further evidence that short-term grammar learning is enhanced in the context of non-uniform musical scales [15], suggesting that their striking prevalence across musical cultures [10] may stem from deeper cognitive principles of learning [32; 33].

Building on the same design, the present study establishes that these facilitating effects originate from enhanced neural encoding of melodies derived from non-uniform scales. Non-uniform scales, such as the pentatonic scale and Western diatonic scales, are characterized by a specific positioning of the tones along the octave that create a unique set of relationships among them. This organization promotes the perception of a tonal hierarchy, a key aspect for melodic processing [32; 33]. In contrast, uniform scales, which are very rarely observed [10; 34], are characterized by an even distribution of tones around the octave, resulting in an entirely non-specified tonal space [15].

Here, statistical learning of melodies was investigated in the context of a uniform and non-uniform scale, schematically represented in Figure 1.A. To generate statistical properties governing melodic syntax, an artificial first-order grammar (Figure 1.B) was used to generate melodies. First, in an exposure phase, listeners were presented with a corpus of 100 *reference* melodies generated from the original grammar. In the following test phase, half of the presented melodies were from the original grammar (i.e. *reference* melodies), while the other half contained wrong (out-of-grammar) transitions that were inserted to induce syntactic violations in the melodies (i.e. *alternative* melodies). Listeners had to report whether each of the melodies sounded familiar with respect to what they heard in the exposure phase. This procedure was repeated two times, once for each scale condition. Details of the experimental setup, subjects, and stimuli are provided in **Methods** section.

**Figure 1:**
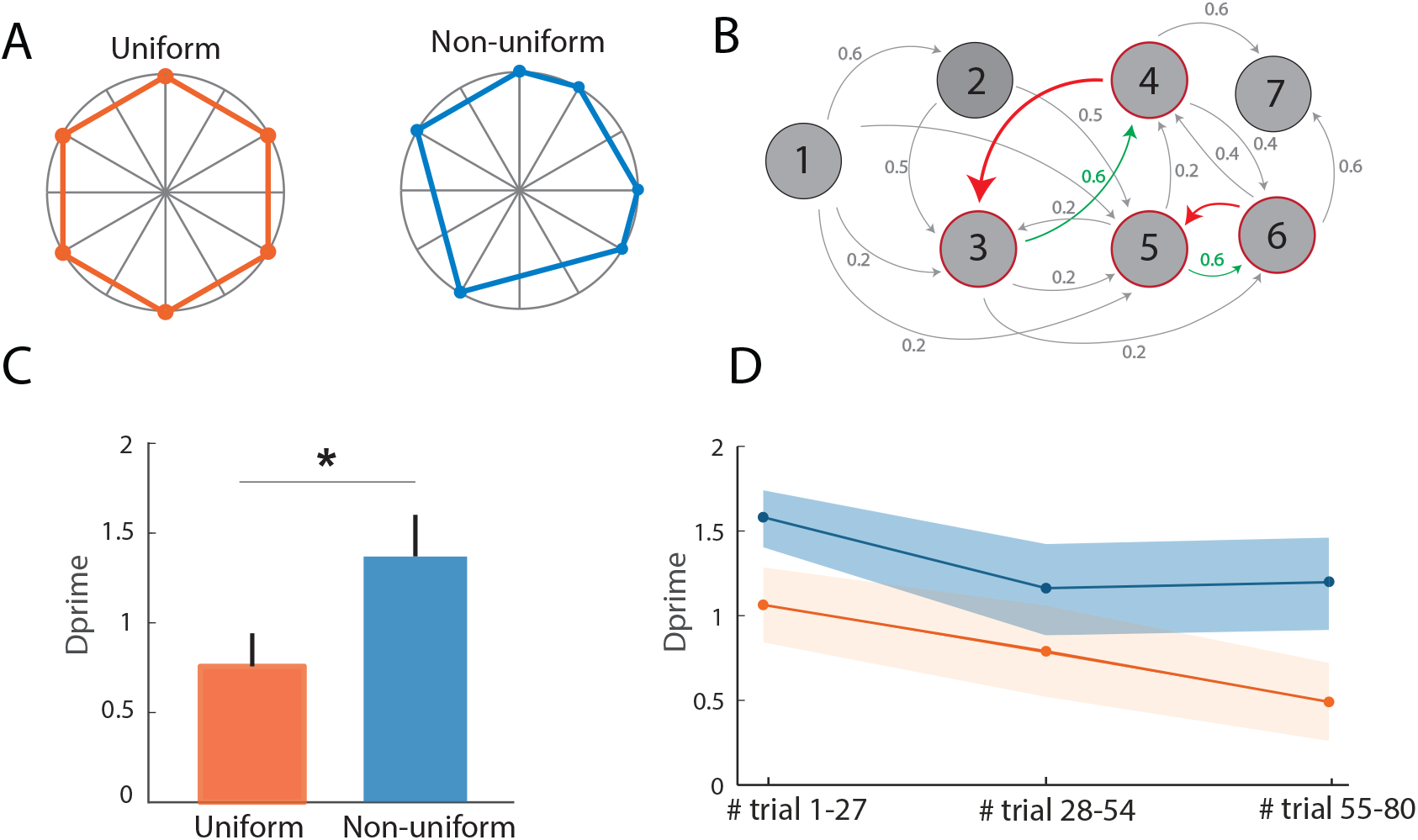
Method and Behavioral data. **(A)** Schematic representation of the uniform (red) and non-uniform (blue) scales as circular diagrams. **(B)** The first-order Markov-chain grammar used to generate melodies from the uniform and non-uniform scales. Nodes represent scale notes; gray and green arrows represent the transitions between nodes used to generate exposure and reference melodies; and red arrows represent two possible examples of “incorrect” transitions used to generate half of the test melodies (i.e. the alternative melodies). **(C)** *d*^*′*^ values averaged across participants by symmetry condition: uniform (red) and non-uniform (blue). Error bars correspond to standard error. **(D)** To account for the drift in performance during the test session, *d*^*′*^ values are averaged across participants for three different trial groups: trials 1-27, trials 28-54 and trials 55-80.

### Behavioral results reveal learning benefits for non-uniform scales

Following the exposure phase, listeners were presented with either a reference or an alternative (i.e. containing incorrect transitions) melodies. As a way of probing syntactic learning, participants were asked to report whether or not the melody they heard in each trial sounded familiar with respect to what they heard in the exposure phase [15; 35]. Mean *d*^*′*^ values for both scales were computed to assess performance and are shown in Figure Figure 1.C. Performance was significantly higher when melodies were generated with the non-uniform scale (paired *t*-test, *p* = 0.023). Over the course of the test phase, listeners were presented with more and more incorrect transitions in the melodies, which could potentially alter the representation of the grammar they acquired throughout the exposure. To account for this expected drift in performance over time, mean *d*^*′*^ values were averaged across listeners for evenly divided sets of trials over time (first set: trials 1-27, middle set: trials 28-55 and last set: trials 56-80, Figure 1.D). For both scales, a significant drift in performance over time was observed (two-way repeated-measure ANOVA: *F* (14, 2) = 4.75, *p* = 0.017).

### Neural correlates of melodic learning are modulated by scale conditions

To investigate the neural basis of this behavioral effect, we sought to probe the difference in grammar encoding for uniform and non-uniform scales using multivariate pattern analysis applied to neurophysiological data [36; 37; 38], a method that has advanced the understanding of spatio-temporal neural activations related to stimulus processing [39; 40; 41; 42; 43]. The idea is to train a set of linear classifiers to classify sets of neural data collected under different conditions. This provides insight into how the topographical maps collected with EEG sensors display a pattern of activity can discriminate between stimulus features [36; 44; 45; 46], and also reveals how this discrimination evolves over time [36; 47; 48; 46; 41].

Here, we trained a set of linear classifiers to discriminate between the reference melodies (i.e. those generated with the same grammar as during the exposure phase) and the alternative melodies (i.e. those containing syntactic violations). We first used the EEG signal collected over the entire melody to probe the temporal dynamics of neural representation across the whole duration of the melodies. The method consisted in training a set of independent logistic regressors to discriminate signals using all 64 sensors as detailed in the **Methods** section. The classifiers were trained to linearly separate the two melodic conditions (i.e. reference versus alternative) based on the EEG topographic maps at different time points. In addition, we probed the time generalization dynamics [36]: the classifiers were trained on the EEG response of 6 seconds of melodies at each time point *t* and tested at time *t*^*′*^, where *t* and *t*^*′*^ were different time points sampled over the time-window that spanned the entire duration of the melodies (0 to 6 seconds). In order to investigate neural representation of reference and alternative melodies from uniform and non-uniform scales, the analysis was conducted separately using the data-sets collected during each test phase of both scale conditions.

Figure 2.A illustrates the time generalization dynamics of decoders’ performance for the melodies generated from the uniform (left) and non-uniform (right) scales. The classifiers’ scores were significantly above chance level for the non-uniform scale (*p*_*min*_ *<* 0.05) between 3 and 5 seconds after onset of the first tone. This demonstrates that listeners could learn the unfamiliar and artificial musical grammar from melodies generated in this scale. The pattern of the significance region (right) suggests a temporally jittered activity due to the small variations in the emergence of the effects across subjects [36]. Conversely, the classifiers could not discriminate between the reference and alternative melodies when melodies were generated using the uniform scales (*p*_*min*_ *>* 0.05). To simplify the visualization, we directly compared the diagonal scores in non-uniform and uniform scales (i.e., the scores for which testing and training data were synchronized); we observed a statistical difference between the classifier performances, in which the area under the receiver operating characteristic curve (AUC) was significantly larger for the non-uniform scale at around [3-5] seconds following the melody onsets (Figure 2.B).

**Figure 2:**
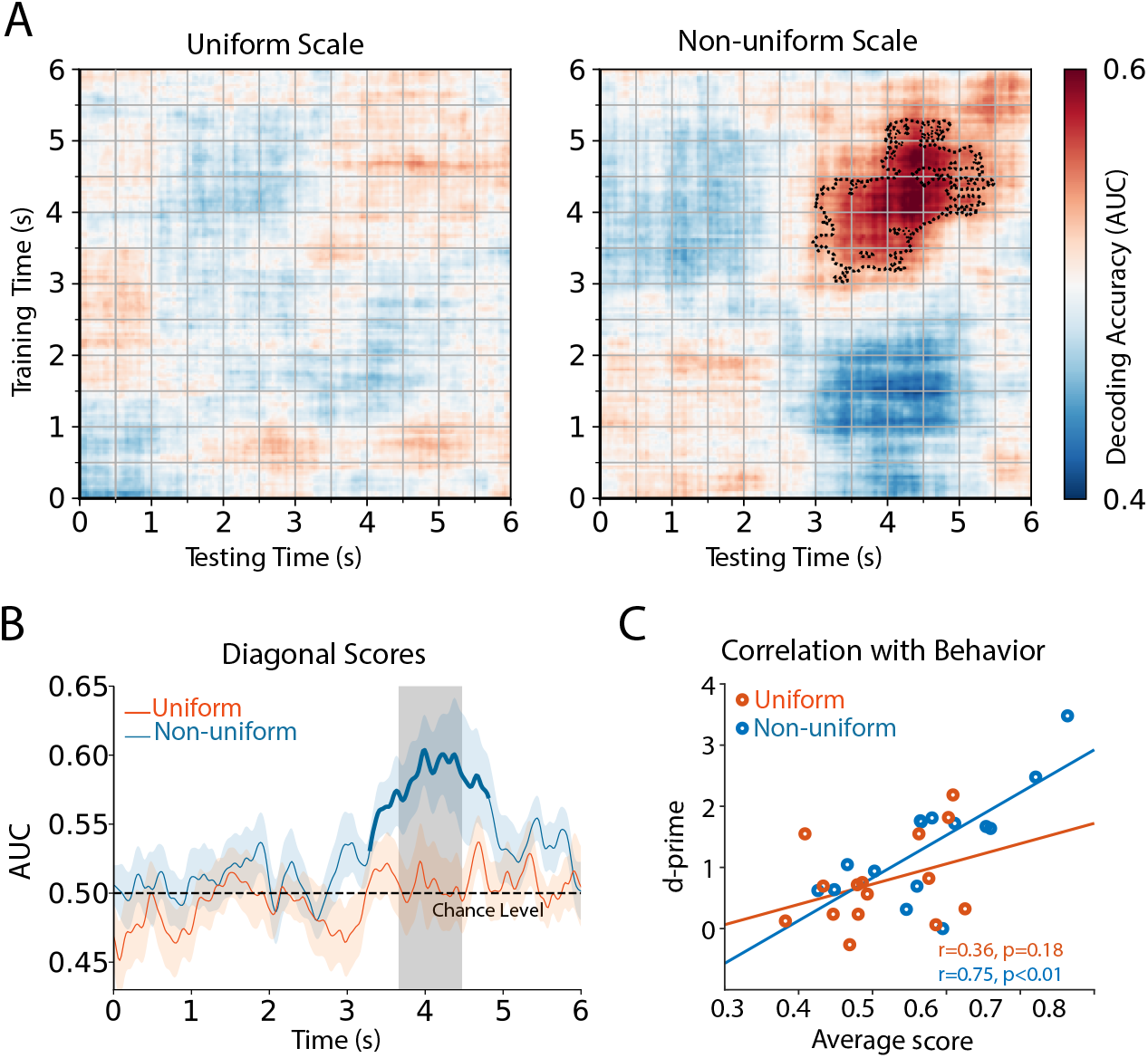
Decoding alternative versus reference melodies. **(A)** Decoding performance for *uniform* (left) and *asymmetric* (right) scales. Classifiers were trained and tested separately at each time in a 6-second time window following the onset of melodies. Cluster-corrected significance is contoured with a dashed line. The classifier scores were significantly above chance level only for the asymmetric scale (*p <* 0.05). **(B)** Decoding performance for the same training and testing time points, which is equal to the diagonal scores in part **A**. The bold curve marks time points where predictions were significantly above chance level (*p <* 0.05). The difference between the scores in uniform and non-uniform scales was significant for the gray bar. **(C)** The average decoding scores for the significant times (bold curves in part B) were correlated with the corresponding behavioral performance, across subjects. The decoding scores and *d*^*′*^ values are significantly correlated only for the non-uniform scale.

**Figure 3:**
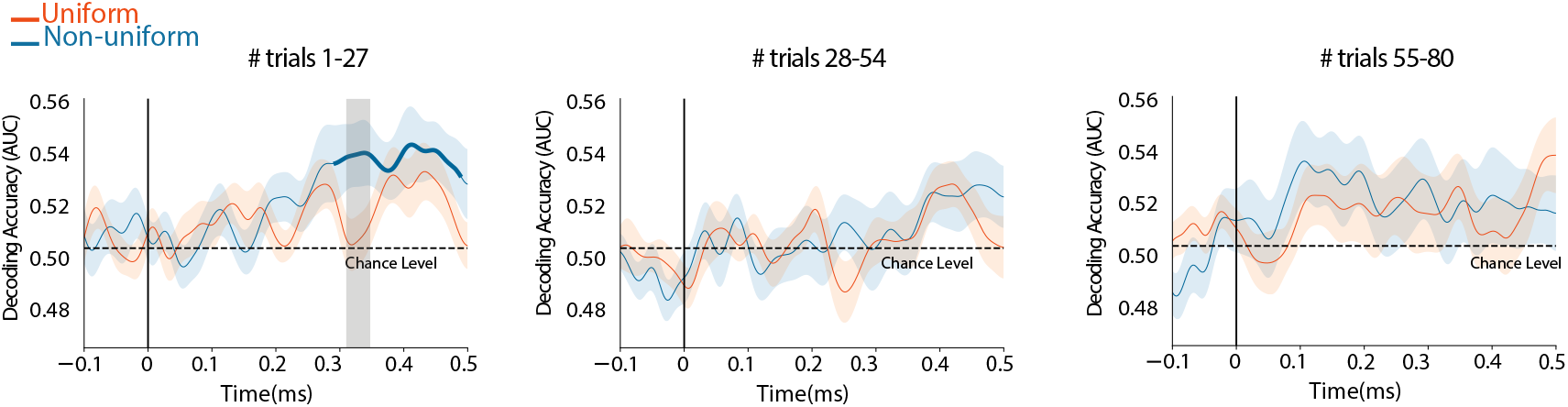
Decoding the *correct* versus *incorrect* transitions. Decoding performance for *uniform* (orange) and *asymmetric* (blue) scales in three separate time windows. Time windows 1, 2, and 3 referred to the first, second, and third portions of the trials. Cluster-corrected significance is marked with a bold line. The classifier scores were significantly above the chance level only for the asymmetric scale (*p <* 0.05) in the the first time window. The difference between the scores in uniform and asymmetric scales was significant for the gray bar.

To directly test whether the enhanced decoding accuracy was related to behavioral performances, we computed the correlation between the classifier accuracy and the *d*^*′*^ values at the individual level.

For the decoding scores, we averaged over the time window in which there were scores significantly above chance level (3-5 seconds) and found a significant correlation for the non-uniform scale (*r* = 0.75, *p <* 0.01) as seen in Figure 2.C. These results suggest a direct link between enhanced performance of the classifiers and behavioral performance on the task.

### The Neural Encoding of correct *vs* incorrect note transitions within the two scale contexts

In the previous analysis, the decoding accuracy rose above chance level at about 3 seconds after melody onsets, which is on average the time when the first non-syntactic notes from the alternative grammar are played, as detailed in the **Methods** section. To investigate more directly the encoding of note transitions in the two scale contexts, we applied a set of linear classifiers to decode the neural data specifically collected during correct and incorrect transitions. Informed by the drift in behavioral performance for both scales (see Figure 1.D), we divided the neural data into the corresponding time periods prior to training classifiers. As in the earlier findings, this analysis revealed that decoding scores were significantly above chance only for the non-uniform scale, starting at 350 ms after the onset of the tones (*p <* 0.05) and only for the first set of trials, due to the drift in performance over time. By contrast, the classifier accuracy remained near chance levels for all the trial sets of data collected under the uniform scale condition.

### Scale-dependent syntactic processing revealed by the evoked responses of musical notes

In the previous analysis, scale effects were probed by comparing differences in decoding accuracy between reference versus alternative melodies, and by examining the topographic maps of EEG signal collected for melodies under both scale conditions. A more direct evaluation of the neural responses elicited by different scales is to compute the evoked responses time-locked to musical events in the melodies. “evoked responses” refer to the average from many repetitions of the neural response elicited by a specific stimulus event. This averaging removes noise and extracts specific negative or positive amplitude peaks directly related to stimulus processing. Here, we sought to investigate the evoked responses that are time-locked to correct and incorrect transitions, and hence gain insights into the type of processes at play under the two scale conditions. Because of the relatively small number of repetitions applied in each condition, we first denoised the data using **D**enoised **S**ource **S**eparation (DSS) [49] as detailed in the Methods section.

This algorithm consists of selecting components of the neural signals that are most repeatable across stimulus repetitions and therefore likely to reflect stimulus processing instead of noise. The components obtained are then projected back onto the sensor space, resulting in a clean EEG signal for each of the 64 channels. We used the DSS-denoised signal from Cz (the electrode situated on the mid-line sagittal plane center, as typically done for the analysis of auditory components) and time-locked between -100-400 ms to notes in the alternative melodies with incorrect transitions, i.e. potentially exhibiting a syntactic violation to the exposure grammar. As seen earlier, we took into account the drift over time in performance by dividing the neural data into the corresponding time periods. The resulting DSS-denoised evoked responses are plotted in Figure 4.A. A bootstrap re-sampling conducted on evoked responses from correct and incorrect transitions revealed that the two signals were significantly different *only* for the non-uniform scale and in the first time-window (see **Methods** section for details on the statistical analysis).

**Figure 4:**
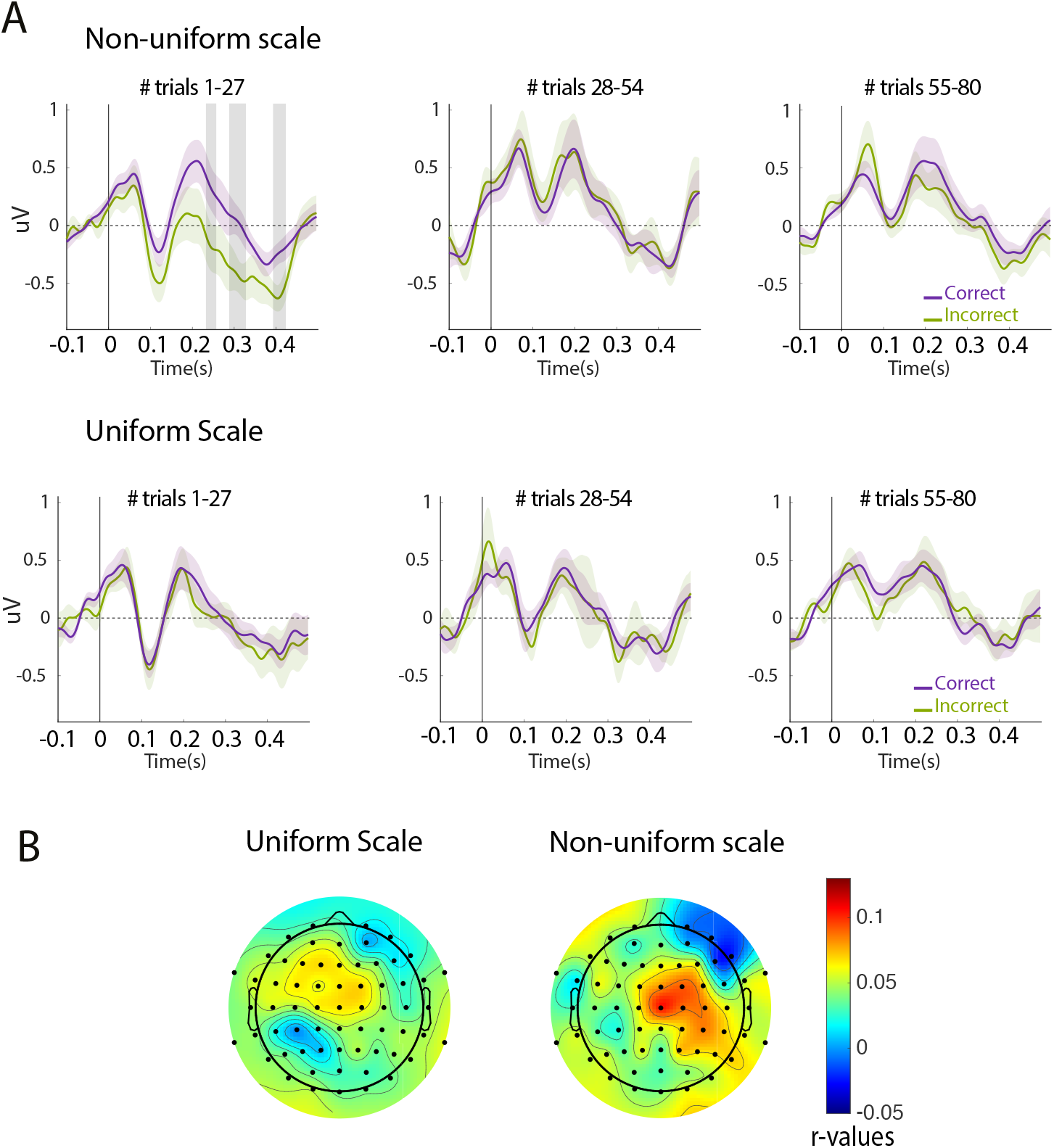
Temporal and topographic markers of syntactic processing. **(A)** Evoked responses at channel Cz for *correct* and *incorrect* transitions. Comparison between the evoked responses due to the *correct* transitions with *incorrect* transitions for three separate sets of trials. Time windows 1, 2, and 3 refer to the first, second, and third portions of the trials. There were significant differences between the *correct* and *incorrect* transitions only in *time window 1* for the asymmetric scales. **(B)** Topography of difference in grammar encoding. Non-uniform - Uniform *r*-values obtained from the TRF of probability of notes for all melodies (reference and alternative) are first subtracted from a null model (100 permutations).

The evoked response of incorrect transitions for the non-uniform scale revealed a larger negative component between 200 and 400 ms after note onset. This late response elicited by an ungrammatical event is strongly evocative of the well-known ERAN component associated with syntactic processing in the context of music [50; 51; 52; 53; 54] and also referred as the “musical MMN component” [55].

### Global temporal and topographic characterization of the neural correlates of different scales

Finally, we examined the general sensor topography of grammar encoding under different scale conditions. For that, a Temporal Response Function (TRF) analysis [56] was conducted on the neural data collected for the uniform and non-uniform scales. The TRF is a decoding technique used to account for the neural encoding of continuous stimulus features such as envelope, semantics, or phoneme for speech [57; 58; 59; 60] and envelope and syntax for music [61; 62]. We used the -log probability of note transitions between the notes estimated for each melody (i.e., the syntactic structure) based on the reference grammar, which is very similar to the *surprisal signal* [63], a computation that provides good measures for interpretation of perceptual data [64; 65]. Regressing surprisal from neural signal using TRF has been shown to yield significant cortical tracking of syntactic structure during melodic processing [62; 66; 67].

The TRF is essentially a kernel that describes the linear mapping of the stimulus into the neural response using ridge regression. The kernel is fit to minimize the mean-squared error between the actual neural response and the predicted neural response. The encoding index is then assessed using a cross-validated evaluation using *r* Pearson’s correlations between the predicted and actual neural responses for each individual channel. Details of the TRF analysis are provided in the **Methods**. section. In brief, this analysis provides an index of how well the grammatical structure of the melodies is represented at each EEG sensor, thus revealing the general topography of the neural processing of syntax under both scale conditions.

A TRF was computed for each data set and each condition and trained on trials in the test phase. Then, Pearson’s correlations for only the first third of trials were averaged over participants and plotted in Figure 4.B. The topographies reveal that for both scale conditions, the electrodes tracking the syntactic information are situated in the central region. They also reveal a significant effect of scale conditions (cluster-based permutation test, 2,000 iterations *p <* 0.01) observed on the centro-lateral electrodes. This is consistent with earlier findings of syntactic processing in the context of music listening that report higher correlation with surprisal in these electrode regions [62; 66; 67], which further confirms that the main effect of scale condition is driven by differences in the grammatical learning of the unfamiliar melodic corpus [15].

## Conclusion

Music is a universal feature of human societies but displays tremendous variability in its rhythmic, timbral, and pitch structure [4; 5; 10]. Yet converging findings from large datasets of musical production suggest that the musical world exhibits some quasi-universal structural traits [34; 68; 10] that could reveal common underlying cognitive properties in the processing of musical signals. In particular, musical scales with a non-uniform (asymmetric) tonal structure (i.e. note sequences separated by intervals of varying sizes) are far more prevalent than uniform ones across musical systems [34; 10]. This motivates the hypothesis that these differences stem from the cognitive benefits of asymmetric scales in melodic learning [32; 33; 69].

Recent findings have further supported this hypothesis. For instance, a recent behavioral study highlighted the benefit conveyed by non-uniform scales for the learning of unfamiliar syntactic rules [15]. Non-uniform scales were found to enhance performance on a syntactic violation task within the context of familiar and unfamiliar musical systems. The benefit may originate from an enhanced internal representation of the tonal space in which relations among tones are better delineated by a non-uniform scale [32]. This study explored the neural correlates and origins of this preference by associating the behavioral effects of enhanced performance with simultaneous measurements of the neural responses while subjects learned melodies generated within uniform and non-uniform scales [15].

The behavioral and neural data were collected during a test phase in which, after only a short exposure to reference melodies, listeners’ ability to learn an artificial musical grammar was probed through their performance in detecting syntactic errors in alternative versus reference melodies. The results confirmed that the superior learning for melodies generated with the non-uniform scale—as indicated by higher *d*^*′*^ values—was paralleled by an enhanced neural encoding of that grammar. The latter findings were assessed by linear classifiers that better discriminated the topographical maps of EEG activation when the melodies were generated from the non-uniform scale. Finally, evoked responses for incorrect and correct transitions were also modulated by the scale condition: in the case of non-uniform scales, a late negative component evocative of the ERAN [55; 53; 54] was observed for incorrect transitions embedded in alternative melodies of the non-uniform scale.

In summary, we have demonstrated that the twos type of melodies, alternative and reference, were better represented in the neural data when they were derived from the non-uniform scale, and that this was correlated with behavioral performance at the individual level. Further analyses of temporal and spatial characteristics of the EEG responses confirmed that the effect of scale type was driven by a more efficient syntactic processing in the context of the non-uniform scale.

Humans spontaneously seek musical experiences for pleasure [70; 71; 72], arguably in search of social bonding that reinforces human survival and reproduction [73; 74; 75; 76; 77; 78; 79; 80]. This highlights the relevance of music production and learning for cultural evolutionary theory [81]. Numerous and converging studies demonstrate that humans learn the musical regularities of their own cultural background [18; 19; 20; 23] from a very early age [82; 83], throughout life [26; 27; 28], and in the absence of explicit instructions, much like they learn their mother tongue [84]. In this perspective, musical features could be selected through evolutionary processes to enhance music learning and thus favor common pleasurable experiences and social bonding [74; 85; 81].

This study provides evidence of the facilitating effects in learning unfamiliar music in the context of a scale structure that is most prevalent across musical structures. This behavioral effect was paralleled by neural findings showing better neural encoding of melodies, most likely related to enhanced syntactic processing. Altogether, these results bring strong new evidence that cross-cultural universals in the music domain reveal cognitive principles of auditory processing.

## Competing interests

Authors declare no competing interest.

## Author Contributions

CP, MR, MF and SS: conceptualization. CP and MR: collection of data. CP and MR: analysis of data. CP, MR, MF and SS: writing and editing.

## Acknowledgments

We thank David Poeppel and Omri Raccah for their thorough comments on the manuscript.

## Funding

This study was supported by an Advanced European Research Council grant (NEUME, 787836) and Air Force Office of Scientific Research and National Science Foundation grants to S.A.S.

## Method

### Participants

Sixteen adult participants with self-reported normal-hearing participated in this study, conducted at the University of Maryland. One participant was removed from the analysis for not keeping the earphones in place during the experimental procedure. Among the fifteen remaining participants (9 females, mean age = 25 years, *SD* = 8), two had five or more years of formal musical training and all were still engaged in daily musical practice. All participants were given course credits or monetary compensation for their participation. The experimental procedures were approved by the University of Maryland Institutional Review boards. Written informed consent was obtained from each subject before the experiment.

### Scale

Participants were presented with melodies generated from hexatonic scales in the two following structure conditions: uniform and non-uniform. Each scale was composed of six tones in 12-tone equal temperament (12-TET). The two scale conditions were obtained by positioning 12-TET tones in a manner that conformed to the different intervallic structural properties, as illustrated in Figure 1.A. The uniform scale was composed of intervals (i.e., space in between the pitch of subsequent tones) of equal sizes. In contrast, the non-uniform scale was composed of intervals of different sizes so that each tone had a unique set of intervallic relations with all of the others tones when moving from one tone to another in the same direction (clockwise/counter-clockwise or up/down) across the octave span.

### Grammar

Melodies were composed of the tones within a given scale, and their construction was determined by a first-order Markov chain inspired by Rohrmeier et al. [86]. Since two scale types were used (hexatonic, 12-TET), two different grammars were used as well, each including all the tones in the scales. However, the complexity of both grammars was kept constant with respect to the transition probabilities that were used to generate the melodies. Schematic representations of the grammar are shown in Figure 1.B. Each node corresponds to a tone in the scale. The correspondence between nodes and notes was randomized for each participant and for each structure condition. Arrows connecting nodes determine the permissible transitions between notes, along with the probability of transition. The “reference” version of the grammar was determined for each listener prior to the exposure phase. The “alternative” version of the grammar was obtained by switching nodes 3-4 and 5-6, which introduced 10 possible wrong transitions. Melodies generated with the alternative grammar contained a set of three transitions between tones that were never part of the melodies generated with the reference grammar.

### Melodies

All melodies were composed of 500 ms sine tones to which a tapered-cosine (Tukey) window was applied. Tones were not separated by a silence interval. During the exposure phase, 100 melodies were generated in real time using the grammar structure and the pitches of tones defined by each scale. During the exposure phase, melodies were produced using the current structure condition (uniform or non-uniform) and the reference version of the grammar. During the test phase, half of the melodies were produced the same way using the reference grammar and half of the melodies were produced the same way but using the alternative grammar. Forty reference and 40 alternative melodies were presented in a random order during the test phase. All melodies were constrained so that they did not exceed 15 tones and had to reach the final note, as defined by the grammar.

### Procedure

The experiment was divided into two parts, each part corresponding to a structure condition in which order of testing was randomized across participants. During each part, listeners had to first complete an exposure phase during which they listened to 100 melodies. During this phase, melodies were generated in real time with the designated scale and grammar. Only the correct version of the grammar was used to generate the exposure melodies. Throughout this phase, listeners had to simply click a mouse to play the next melody. Immediately following the exposure phase, participants completed a test phase during which 80 melodies were generated on the fly; half of them were generated using the reference version of the grammar and the other half with the alternative version. After each melody, participants had to report whether this melody sounded familiar or unfamiliar, with respect to what they just were exposed to in the previous phase. Participants were tested individually in an EEG testing booth. Audio files of the stimuli were encoded at 16-bit resolution and 44.1 kHz sampling rate and presented via Etymotics Research ER-2 earphones. The stimuli were presented at a comfortable loudness level above 60 dB SPL (A-weighted). Instructions were displayed on a computer screen and participants’ responses were collected with a keyboard and mouse. Informal debriefing with participants indicated that both scales were perceived as equally unfamiliar and no formal ratings of familiarity were collected after each session.

### Data Acquisition

Electroencephalogram (EEG) data were recorded using a 64-channel system (ActiCap, BrainProducts) at a sampling rate of 500 Hz with one ground electrode and re-referenced to the average. We used a default fabric head-cap that holds the electrodes (EasyCap, Equidistant layout).

### EEG Prepossessing

EEG data was first mean-centered to perform zero-order detrending. We detected bad channels as exhibiting amplitude above 3 standard deviation from the channel average. Selected bad channels were then interpolated using a weighed sum of neighboring channels’ signal. To avoid artifacts caused by low-pass filtering, we subtracted from each channel its slow varying trend by robust-fitting a 30*th*-order polynomial [87]. We then applied an anti-aliasing low-pass Butterworth (IIR) 4-order filter with a 40 Hz cut-off and down-sampled the resulting data to 100 Hz. Using a time-shift PCA, eye-blink artifacts were isolated and projected out using data collected by the HEoG and VEoG channels [88]. Finally, the EEG data was re-referenced again by subtracting the robust mean (as defined in [87]) before it was epoched using the triggers sent at the beginning of each trial. Bad epochs were selected based on an amplitude above 3 standard deviation and discarded for the analysis.

### Decoding

To evaluate the separability of neural traces elicited by melodies from the two scales, we trained a set of logistic regression classifiers on the preprocessed EEG data (e.g. not DSS-denoised). At each time point *t* we used the matrix of observations *X*_*t*_ *∈ R*^*N×*64^, for *N* samples of all 64 electrodes to predict the labels *y*_*t*_*′ ∈ {*0, 1*}*^*N*^. Here, the labels corresponded to the two grammar conditions (alternative vs. reference melodies) or the state of transitions (correct vs. incorrect), or the probability of transitions (low probability vs. high probability transitions). This was repeated for every time point *t*^*′*^ of each epoch. This analysis was conducted two times: on the epochs collected from the uniform scale conditions and the non-uniform scale condition. For each subject, we trained the decoders on EEG signals at each time points of the melodies (from onset to 6 s after onset). Therefore, the decoder at each time point learns to predict the grammar conditions (alternative vs. reference) using the topography of the EEG samples for this time point. Additionally, temporal generalization analysis was conducted to capture the dynamics of topographical patterns of EEG signal over time (for more details on that, see [36]). To achieve that, we systematically evaluated each classifier from each time point to all other time points. Concretely, this means that a classifier trained to separate labels at a given time point is then used to predict the labels at all other time points.

To validate the classifier’s performance, we used 5-fold cross-validation. This means that for each individual data set, over 5 iterations, the trained classifier was used to predict labels on a fifth portion of unseen data. The area under the receiver operating characteristic curve (AUC) was used to quantify the classifier’s performance. We implemented this decoding analysis using sci-kitlearn [89] and MNE [90] libraries in python 3.6.

### Temporal Response Function

To evaluate the different topographical mapping in melodic encoding between the two scale structures, we used a brain decoding method based on Temporal Response Functions (TRF) [56]. The TRF is based on a class of linear time-invariant models that describes the linear transformation of stimuli features to the neural signal (EEG) by its impulse response after ridge regression. Unlike an evoked response, the response function obtained reflects a modeled neural response to a *specific* set of features (when an ERP represents the grand average to the whole stimulus). More precisely, the TRF optimally describes the mapping between a given set of features of a sensory input *s*(*t*) and the neural response *r*(*t*) collected from each channel *n* of the neural signal such as defined in Eq. 1:

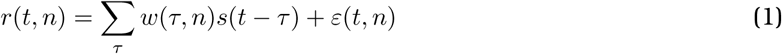

Where *τ* is the specific range of lags for which the response at time *t* is described (here, [-100 - 500] ms) and *ε*(*t*) is the residual error at each channel *n* not explained by the model. The TRF *w*(*τ, n*) is estimated by minimizing the mean-squared error between the actual neural response and the one predicted by the convolution *w*(*τ, n*) *\* s*(*t − τ*). The model is optimized using ridge regression and assuming a certain degree of regularization to prevent over-fitting. This regularization parameter is optimized in the [10^*−*3^, 10^3^] interval, using logarithmic steps, and for each individual data set. To evaluate the performance of the model, a cross-validated via leave-one-out evaluation using Pearson’s correlations between the predicted and actual neural responses is conducted. The resulting topographical map indicates the strength of stimulus feature encoding at each EEG channel. Prior to conducting the TRF analysis, a visual inspection of trials was done to remove noisy portions of the data. Additionally, disparate external noise sources were removed by conducting an ICA and removing components which topography indicated signal from an external source.

### Denoised evoked responses

For the the evoked response analysis, a specific denoising algorithm called Denoised Source Separation (DSS) was applied (for detailed explanation see [49]). In a nutshell, DSS isolates components of signals that are mostly repeated across repetitions of trials, so as to keep the relevant signal (e.g. one that reflects stimuli properties) and to remove signal resulting from noise. In the present study, the Denoised Source Separation (DSS) filter’s output was the weighted sum of the signals from the 64 EEG electrodes, in which the weights were optimized to extract the repeated neural activities across trials. This transformation yielded to 64 uncorrelated brain source activities (e.g., DSS components) which were ordered by a repeatability score. Since the trials were not exactly identical between repetition, we selected only the first 5 most repeatable DSS components and projected them back in the sensor space to obtain cleaned signals. Finally, we used the obtained denoised Cz electrode (placed on the mid-line sagittal plane center) for the evoked response analysis.

### Statistical Analysis

#### Classifiers

Statistical analysis for the classifiers was performed with a one-sample *t*-test with random-effect Monte-Carlo cluster statistics for multiple comparison correction using the default parameters of the MNE spatio_temporal_cluster_1samp_test function [91]. Error bars in all figures represents ±SEM (standard error of the mean).

#### Analysis of evoked responses

To compare the averaged evoked response between conditions, we performed bootstrap resampling in order to estimate the standard deviation (SD) of the difference between the type of transitions (correct vs. incorrect). Significance levels were set for difference between the two signals above 2*×* estimated SD. In Figure 4.A, error bars represents ±SEM (standard error of the mean).

#### Topography

In order to assess the different topographies for both scale conditions, we conducted a one-sample, cluster-based permutation test using *r*-values as input [92]. In this analysis, multiple *t*-tests are computed for each electrode. Then, best on clusters of electrodes in which the response significantly differs from zero are identified. These clusters are then formed over space by grouping electrodes that have significant initial *t*-test values. The sum of all *t*-scores within each cluster provides a cluster-level *t*-score (mass *t*-score). A permutation approach is then used to control for Type I errors (2,000 iterations) in order to build a data-driven null hypothesis distribution. The significance of a cluster is determined by whether it falls in the highest 5th percentile of the corresponding distribution (*α* = 0.05).

## Notes

### Competing Interest Statement

The authors have declared no competing interest.

